# CellProfiler Analyst: interactive data exploration, analysis, and classification of large biological image sets

**DOI:** 10.1101/057976

**Authors:** D. Dao, A. N. Fraser, J. Hung, V. Ljosa, S. Singh, A. E. Carpenter

## Abstract

**Summary:** CellProfiler Analyst allows the exploration and visualization of image-based data, together with the classification of complex biological phenotypes, via an interactive user interface designed for biologists and data scientists. CellProfiler Analyst 2.0, completely rewritten in Python, builds on these features and adds enhanced supervised machine learning capabilities (in Classifier), as well as visualization tools to overview an experiment (Plate Viewer and Image Gallery).

**Availability:** CellProfiler Analyst 2.0 is free and open source, available at http://www.cellprofiler.org/releases and from GitHub (https://github.com/CellProfiler/CellProfiler-Analyst) under the BSD license. It is available as a packaged application for Mac OS X and Microsoft Windows and can be compiled for Linux. We implemented an automatic build process that supports nightly updates and regular release cycles for the software.

**Supplementary information:** Supplementary Text 1: Manual to CellProfiler Analyst; updated versions are available at <u>CellProfiler.org/CPA</u>

Supplementary Data 1: Benchmarking performance of classifiers in CPA 2.0 versus CPA 1.0

## 1 Introduction

CellProfiler Analyst is open-source software for biological image-based classification, data exploration, and visualization with an interactive user interface designed for biologists and data scientists. Using data from feature extraction software such as CellProfiler (Kamentsky et al. 2011), CellProfiler Analyst offers easy-to-use tools for exploration and mining of image data, which is being generated in ever increasing amounts, particularly in high-content screens (HCS). Its tools can help identify complex and subtle phenotypes, improve quality control, and provide single-cell and population-level information from experiments.

Some distinctive and critical features of CellProfiler Analyst are its user-friendly object-based machine learning interface, its ability to handle the tremendous scale of HCS experiments (millions of cell images), its brushing and gating capabilities that allow observing relationships among different data displays, and its exploration tools which enable interactively viewing connections between cell-level data and well-level data, and among raw images, processed/segmented images, extracted features, and sample metadata. Compared to other commonly-cited open-source biological image classification software like Ilastik (Sommer et al. 2011), CellCognition (Held et al. 2010), and WND-CHARM (Orlov et al. 2008), CellProfiler Analyst has the advantage of containing companion visualization tools, being suitable for high-throughput datasets, having multiple classifier options, and allowing both cell and field-of-view classification. Compared to command-line-based data exploration software like cellHTS (Boutros et al. 2006) and imageHTS (Pau et al. 2013) and the web tool web CellHTS2 (Pelz et al. 2010), CellProfiler Analyst provides interactive object classification and image viewing. Several other software tools (e.g., the HCDC set of modules for KNIME (Berthold et al. 2008) are no longer available/maintained.

Here we present major improvements to CellProfiler Analyst. Since its original publication (Jones et al. 2008), CellProfiler Analyst has been rewritten in Python (vs. its original language, Java) with significant enhancements. While keeping the original functionality allowing researchers to visualize data through histograms, scatter plots, and density plots and to explore and score phenotypes by sequential gating, the key new features include:

- multiple machine learning algorithms that can be trained to identify multiple phenotypes in single cells or whole fields of view, by simple drag and drop
- more efficient handling of large scale, high-dimensional data
- a gallery view to explore images in an experiment, and cells in individual images, and
- a plate layout view to explore aggregated cell measurements or image thumbnails for single or multiple plates.

## 2 New features in CellProfiler Analyst 2.0

### Classifier

CellProfiler Analyst 1.0 allowed researchers to train a single classifier (Gentle Boosting) to recognize a single phenotype (two-class) in individual cell images (rather than whole fields-of-view) (Jones et al. 2009). In CellProfiler Analyst 2.0, Classifier can perform cell and field-of-view-level classification of multiple phenotypes (multi-class) using popular models like Random Forest, SVM, and AdaBoost from the high performance machine learning library scikit-learn (Pedregosa et al. 2011), which yields a ~200-fold improvement in speed (Supplementary Data 1). First, cell-or whole-image samples from the experiment are fetched and sorted by drag and drop into researcher-defined classes, making up the annotated training set. Fetching can be random, based on filters, based on per-class predictions of an already-trained classifier, or based on active learning. The new active learning option speeds annotation by presenting uncertain cases. In addition, researchers can view full images of each sample and drag and drop cells from the image for annotation. Next, a classifier is trained on this set. After training on the annotated set, a model’s performance can be evaluated by cross validation in the form of a confusion matrix and precision, recall, and F1 score per class. The model can then be used to quantify cell phenotypes on a per image and per experiment basis (or images for image classification).

**Fig. 1.**
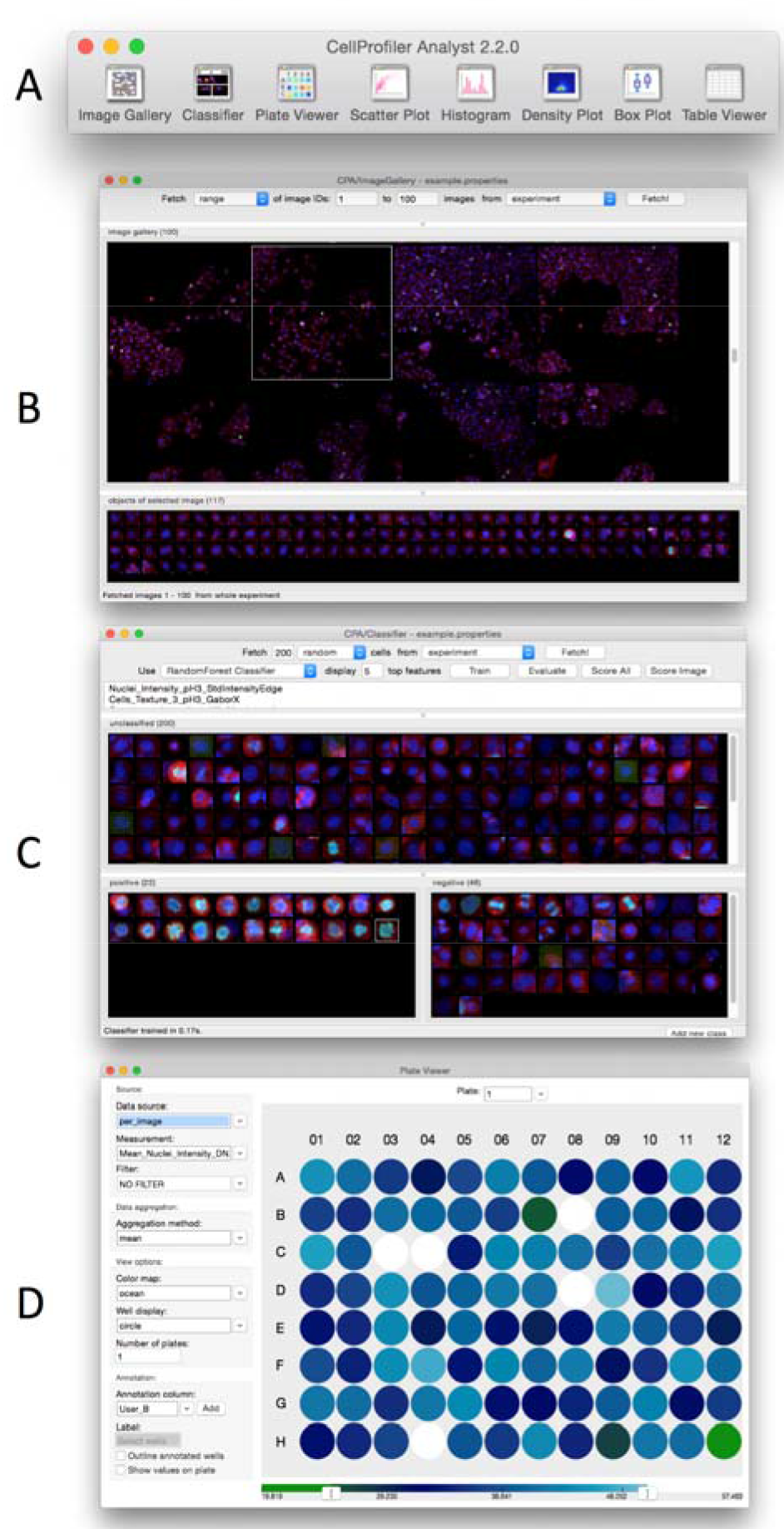
**User Interface of CellProfiler Analyst.** A) Main toolbar; B) Image Gallery; C) Classifier; D) Plate Viewer.

### Image Gallery

CellProfiler Analyst 2.0 offers a convenient new Image Gallery tool, in addition to the existing visualization/exploration tools with standard plotting and gating capabilities in version 1.0 (Jones et al. 2008). Image Gallery provides a convenient gridview allowing an overview of images from the entire experiment. A variety of options are provided to filter images based on experiment-specific metadata, e.g., gene name, compound treatments, etc. multiple filters can be combined to refine the search. Images can be displayed as a custom-sized thumbnail or in full resolution, and the color assigned to each channel in the image can be customized to highlight structures of interest. Individual segmented cells can be viewed for each image, and can be dragged and dropped into the Classifier window.

### Plate Viewer

Many large-scale imaging experiments take place in multi-well plate format. Researchers are often interested in seeing their data overlaid on this format, to check for systematic sample quality issues, or to see results from controls placed in particular locations, at a glance. The new Plate Viewer tool displays aggregated and/or filtered measurements (according to customizable colormaps) or a thumbnail image for each well. Automatically-imported annotations can be viewed, and individual annotations can be manually added or deleted for each well.

### Additional features

Additional features added to CellProfiler Analyst vs. version 1.0 have been described elsewhere, such as Tracer, a tool that complements the object tracking functionality of CellProfiler, including visualization and editing of tracks (Bray and Carpenter 2015), as well as workspaces for saving progress and display settings across sessions (Bray et al. 2012). The website, manual, and tutorials have been redesigned and updated to the new version.

## 3 Future directions

The redesigned CellProfiler Analyst contains useful classification and visualization features in an interactive interface that facilitates data analysis and exploration of biological images. Its code base forms a solid foundation for integrating new classifiers into the tool, potentially including deep learning architectures. We also intend to integrate methods for constructing per-sample “profiles” from raw morphological measurements to support morphological profiling applications (Caicedo, Singh, and Carpenter, in press; Bray et al., Nature Protocols, in press).

## Acknowledgements

The authors thank members of their laboratory for contributing to the development of the software and this manuscript, especially Mark-Anthony Bray, Allen Goodman, Lee Kamentsky, Alison Kozol, David Logan, and Mohammad Rohban.

## Funding

This work has been supported by the National Institutes of Health [R01GM089652 to AEC].

*Conflict of Interest*: none declared.

## References

Bray, Mark-Anthony, and Anne E. Carpenter. 2015. “CellProfiler Tracer: Exploring and Validating High-Throughput, Time-Lapse Microscopy Image Data.” BMC Bioinformatics 16 (November): 368.

Bray, Mark-Anthony, Adam N. Fraser, Thomas P. Hasaka, and Anne E. Carpenter. 2012. “Workflow and Metrics for Image Quality Control in Large-Scale High-Content Screens.” Journal of Biomolecular Screening 17 (2): 266–74.

Bray, Mark-Anthony, Shantanu Singh, Han Han, Chadwick T. Davis, Blake Borgeson, Cathy Hartland, Maria Kost-Alimova, Sigrun M. Gustafsdottir, Christopher C. Gibson, and Anne E. Carpenter. Nature Protocols, in press. “Cell Painting, a High-Content Image-Based Assay for Morphological Profiling Using Multiplexed Fluorescent Dyes.”

Caicedo, Juan C., Shantanu Singh, and Anne E. Carpenter. Curr Opin Biotech, in press. “Applications in Image-Based Profiling of Perturbations.”

Held, Michael, Michael H. A. Schmitz, Bernd Fischer, Thomas Walter, Beate Neumann, Michael H. Olma, Matthias Peter, Jan Ellenberg, and Daniel W. Gerlich. 2010. “CellCognition: Time-Resolved Phenotype Annotation in High-Throughput Live Cell Imaging.” Nature Methods 7 (9): 747–54.

Jones, Thouis R., Anne E. Carpenter, Michael R. Lamprecht, Jason Moffat, Serena J. Silver, Jennifer K. Grenier, Adam B. Castoreno, et al. 2009. “Scoring Diverse Cellular Morphologies in Image-Based Screens with Iterative Feedback and Machine Learning.” Proceedings of the National Academy of Sciences of the United States of America 106 (6): 1826–31.

Jones, Thouis R., In Han Kang, Douglas B. Wheeler, Robert A. Lindquist, Adam Papallo, David M. Sabatini, Polina Golland, and Anne E. Carpenter. 2008. “CellProfiler Analyst: Data Exploration and Analysis Software for Complex Image-Based Screens.” BMC Bioinformatics 9 (November): 482.

Kamentsky, Lee, Thouis R. Jones, Adam Fraser, Mark-Anthony Bray, David J. Logan, Katherine L. Madden, Vebjorn Ljosa, Curtis Rueden, Kevin W. Eliceiri, and Anne E. Carpenter. 2011. “Improved Structure, Function and Compatibility for CellProfiler: Modular High-Throughput Image Analysis Software.” Bioinformatics 27 (8): 1179–80.

Orlov, Nikita, Lior Shamir, Tomasz Macura, Josiah Johnston, D. Mark Eckley, and Ilya G. Goldberg. 2008. “WND-CHARM: Multi-Purpose Image Classification Using Compound Image Transforms.” Pattern Recognition Letters 29 (11): 1684–93.

Pedregosa, Fabian, Gaël Varoquaux, Alexandre Gramfort, Vincent Michel, Bertrand Thirion, Olivier Grisel, Mathieu Blondel, et al. 2011. “Scikit-Learn: Machine Learning in Python.” Journal of Machine Learning Research: JMLR 12 (November). JMLR.org: 2825–30.

Sommer, C., C. Straehle, U. Kothe, and F. A. Hamprecht. 2011. “Ilastik: Interactive Learning and Segmentation Toolkit.” In Biomedical Imaging: From Nano to Macro, 2011 IEEE International Symposium on, 230–33.

